# *Physcomitrium patens* flavodiiron proteins form a redox-dependent heterocomplex

**DOI:** 10.1101/2024.01.29.577648

**Authors:** Claudia Beraldo, Eleonora Traverso, Marco Boschin, Laura Cendron, Tomas Morosinotto, Alessandro Alboresi

## Abstract

Flavodiiron proteins (FLVs) catalyze the reduction of oxygen to water by exploiting electrons from Photosystem I (PSI). In several photosynthetic organisms such as cyanobacteria, green algae, mosses and gymnosperms, FLV-dependent electron flow protects PSI from over-reduction and consequent damage especially under fluctuating light conditions. In this work we investigated biochemical and structural properties of FLVA and FLVB from the model moss *Physcomitrium patens*. The two proteins, expressed and purified from *Escherichia coli*, bind both iron and flavin cofactors and show NAD(P)H oxidase activity as well as oxygen reductase capacities. Moreover, the co-expression of both FLVA and FLVB, coupled to a tandem affinity purification procedure with two different affinity tags, enabled the isolation of the stable and catalytically active FLVA/B hetero multimer protein complex, that has never been isolated and characterized so far. The multimeric organization was shown to be stabilized by inter-subunit disulfide bonds. This investigation provides valuable new information on the biochemical properties of FLVs, with new insights into their *in vivo* role and regulation.

## Introduction

Light is essential for photosynthesis, but when in excess it can damage the photosynthetic apparatus, leading to photo-oxidative stress. As soon as photosynthetic organisms are exposed to saturating light intensity, Photosystem I (PSI) can experience over-reduction due to an excessive flow of electrons reaching its donor side. Under these conditions, PSI can undergo inactivation of its iron-sulfur clusters and photoinhibition. The repair and resynthesis of PSI have significant metabolic costs for the cell, and this process has been demonstrated to occur very slowly over several days (1–3). Essential metabolic pathways, such as the Calvin-Benson cycle and photorespiration, along with cyclic and pseudo-cyclic electron flows support electron flow at the PSI acceptor side and contribute to avoiding over-reduction of PSI (4, 5). Pseudo-cyclic electron flow, also called water-water cycle, involves photosystem II (PSII) that removes electrons from water to produce oxygen (O_2_) and PSI to reduce O_2_ back to water. Pseudo-cyclic electron flow is either sustained by the Mehler reaction or by the activity of flavodiiron proteins (FLVs), enzymes that function as “safety-valves” when linear electron transport though Ferredoxin-NADP^+^-Reductase (FNR) and NADPH pool is saturated (6). FLVs were shown to be active during the induction of photosynthesis in different organisms such as cyanobacteria (7–9), the model microalga *Chlamydomonas reinhardtii* (10), the bryophytes *Marchantia polymorpha* (11) and *Physcomitrium patens* (12) and gymnosperms (13). FLV activity is essential to survive light fluctuations (9, 12).

Photosynthetic FLVs belong to a large family of flavodiiron proteins (usually shortened as FDPs), metalloenzymes that are widespread in Bacteria and Archaea and that catalyze the O_2_ reduction reaction or nitric oxide (NO) reduction reaction. All FDPs have a modular architecture with two conserved core domains (14). At the N-terminus lies a conserved diiron center in a metallo-β-lactamase-like domain enabling electron transfer reactions crucial for O_2_/NO reduction. This is followed by a flavodoxin domain, harboring a flavin mononucleotide (FMN). Apart from the two core domains, different FDPs present various extra modules (14). FLVs found in photosynthetic organisms are characterized by a NAD(P)H:flavin oxidoreductase domain at the C-terminus (15), providing each monomer with a putative electron entry site. The presence of this domain in FLV sequence suggested that NADPH could be the electron donor for the reaction (16, 17). Differently, *in vivo* spectroscopic studies identified reduced ferredoxin as the main electron donor to Flv1/3 for reducing O_2_ to water in cyanobacteria (18). Consistently, in *C. reinhardtii* FLVs compete with [FeFe]-hydrogenases and that could be explained by both proteins using the same electron donor, ferredoxin (19). Crystal structure of Flv1 was recently obtained, but in a version lacking the specific NAD(P)H:flavin oxidoreductase domain (Flv1-ΔFlR) (20), so structural and functional relationship of full length FLVs is still elusive.

Multimeric organization of photosynthetic FLVs is a feature still to be fully clarified. The always coupled FLV genes (21, 22) together with recent experiments carried out in *Synechocystis* and *Physcomitrium patens* suggested that they are active as heterocomplexes. *Synechocystis* strains impaired in either Flv1 or Flv3 lost the FLV ability as PSI electron acceptor and thus showing a higher electron acceptor side limitation of PSI (7, 23) and a strong growth defect if exposed to light fluctuations (24). Also eukaryotes of the green lineage such as green algae, mosses and gymnosperms express two homologous proteins (FLVA and FLVB) (10–12). In photosynthetic red plastid lineage, coral symbionts of the family Symbiodiniaceae have even a polycistronic locus expressing FLVA and FLVB in tandem in a single polypeptide that is cleaved by a post-translational mechanism (25). Also in the moss *Physcomitrium patens* it was shown that the lack of one FLV gene impairs the accumulation also of the other protein, suggesting that they are functional partners, the one stabilizing the other (12). While biochemically FLVA/B were proposed to be heterotetramers based on native gel electrophoresis (26, 27) no *in vitro* studies managed to isolate FLVA/B or Flv1/3 (*Synechocystis* sp. PCC6803) interaction so far. Thus, the aim of this study is to fulfill this lack of information.

In this study, we successfully established the first expression and purification of recombinant flavodiiron proteins from a eukaryotic photosynthetic organism, the moss *Physcomitrium patens*. *P. patens* FLVA and FLVB purified from *Escherichia coli* were shown to bind iron and flavin and displaying NAD(P)H-dependent oxygen reductase activity. We isolated a functional FLVA/B heterocomplex that was shown to be stabilized by inter-molecular disulfide bridges.

## Results

### *P. patens* Flavodiiron proteins sequence and structural prediction analysis

To investigate the properties of plant FLVs, we aligned and analyzed the protein sequences of *P. patens* FLVA (723 *aa*) and FLVB (647 *aa*) (Figure S1). They exhibited 34.68% identity with each other, in line with other members of the same family (28). Based on the amino acid sequence and domain annotation database, we identified the FLVA and FLVB metallo-β-lactamase domain (residues 119-310 for FLVA and 122-310 for FLVB), flavodoxin-like domain (residues 406-545 for FLVA and 340-474 for FLVB) and flavin reductase-like domain (residues 574-723 for FLVA and 498-646 for FLVB) (Figure 1 and Figure S1). FLVA showed specific additional residues (I^313^-K^364^) that are absent in FLVB and that likely form two α-helices connecting the metallo-β-lactamase-like domain and the flavodoxin-like domain. AlphaFold prediction of structures of both enzymes revealed a conserved modular three-domain composition, consistent with the sequence analysis (Figure 1). The metallo-β-lactamase-like domain exhibited the αββα fold characteristic of other metallo-β-lactamase-like proteins. Analysis of the amino acid residues highlighted a classical Asp/Glu/His pattern in FLVB (His_167_-x-Glu_169_-x-Asp_171_His_172_-x61-His_234_-x18-Asp_253_-x56-His_310_) expected to bind the two iron ions of the catalytic site. In FLVA, a diiron binding site was also identified in the protein sequence but with a more basic amino acid composition (His_170_-x-Ser_172_-x-Lys_174_x70-Arg_246_-x18-Lys_265_-x-His_375_). The flavodoxin-like domain displayed the typical αβα topology observed in short-chain flavodoxins. Both FLVA and FLVB possess typical flavin-binding residues in their sequences, hydrophilic residues essential for orthophosphate recruitment, as well as aromatic residues (FLVA Ser_412_-x-Tyr_414_-xx-Asn_416_x80-Ser_496_-Phe_497_-x-Trp_498_, FLVB Ser_346_-x-Tyr_348_-xx-Ser_351_x74-Ser_425_-Phe_426_) (Figure S2A).

**Figure 1.**
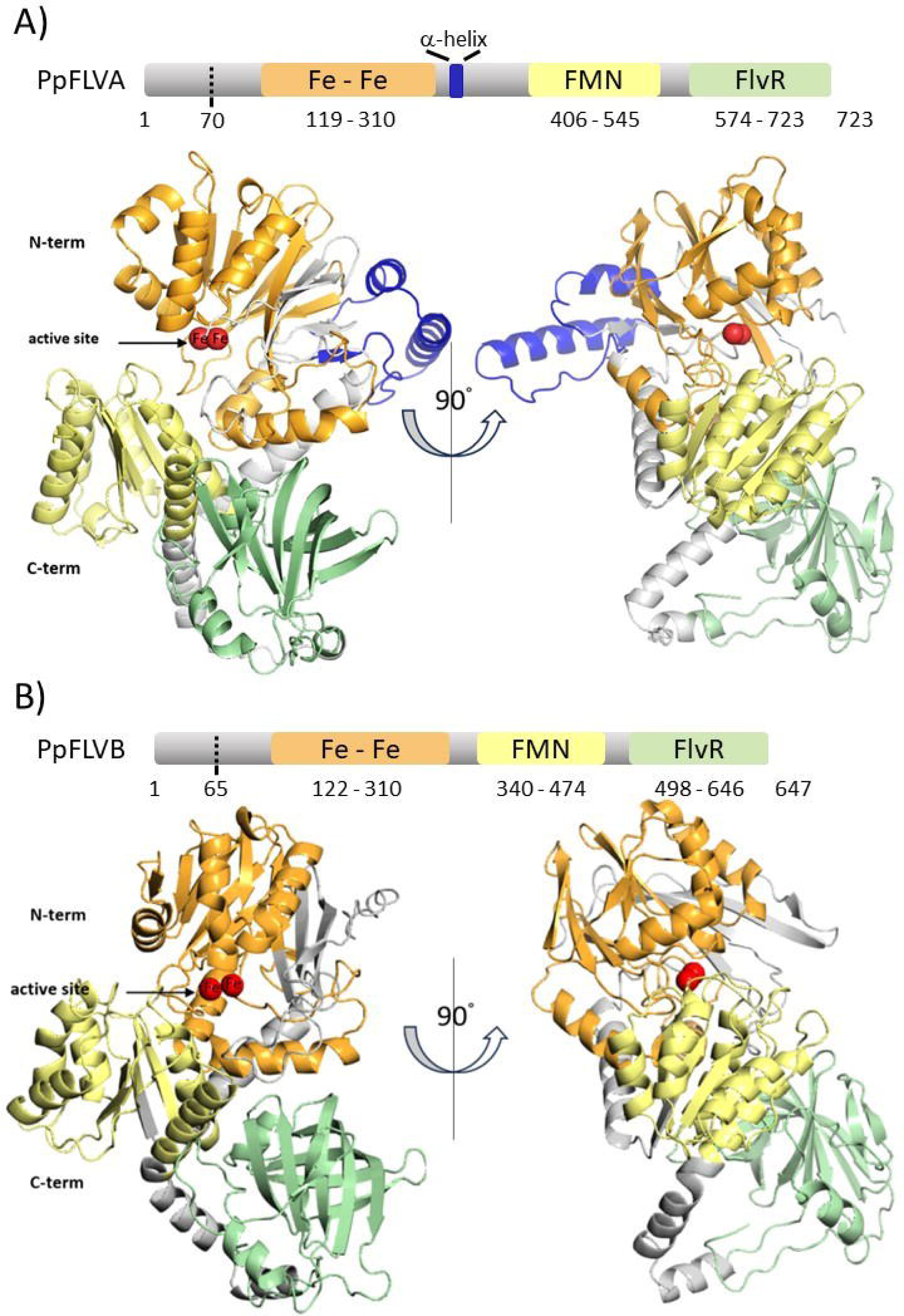
Predicted structure analysis of *Physcomitrium patens* FLVA and FLVB. (A) Schematic representation and AlphaFold model of FLVA (A9RYP3) and (B) FLVB (A9RQ00) structures. Dotted lines indicate the predicted cleavage sites for FLVA (position 70) and FLVB (position 65) chloroplast target peptides. The N-terminal metallo-β-lactamase domain contains a diiron centre (Fe-Fe), the flavodoxin-like domain harbors a flavin mononucleotide (FMN) and the C-terminal domain is a flavin reductase-like domain (FlvR). The three domains are shown respectively in orange, yellow and green. FLVA specific residues (I^313^-K^364^) are shown in blue. Iron atoms (Fe) are represented as red spheres present in the active site.

All this part of the structure is conserved with other FDPs and, as expected, are similar to the partial Flv1-ΔFlR protein from *Synechocystis* (20) that is a homologue of *P. patens* FLVA (22) (Figure S2B). The models provided predictions also for the C-terminal domains of FLVA and FLVB, which are flavin reductase-like typical only of FDPs found in photosynthetic organisms and were absent from any available structure. These C-terminal domains were suggested to be predominantly composed of β-sheets. (Figure 1 and Figure S1).

### Recombinant *P. patens* flavodiiron proteins bind flavin and iron

To characterize *P. patens* FLVs, the recombinant mature FLVA and FLVB were purified by affinity chromatography after heterologous expression in *Escherichia coli*. A StrepII-tag and a 6xHis-tag were respectively fused at the N-terminal part of mature FLVA and FLVB deprived of the chloroplast target peptide. An excess of riboflavin and ammonium iron (II) sulfate was added to the bacterial culture to guarantee the optimal incorporation of the co-factors. StrepII-Tag-FLVA and 6xHisTag-FLVB were expressed independently, and their expression and purification were validated through SDS-page and immunoblot analysis utilizing either antibodies targeting FLVA and Strep-Tag (Figure 2A, Figure S3) or antibodies against FLVB and His-Tag (Figure 2B, Figure S3).

**Figure 2.**
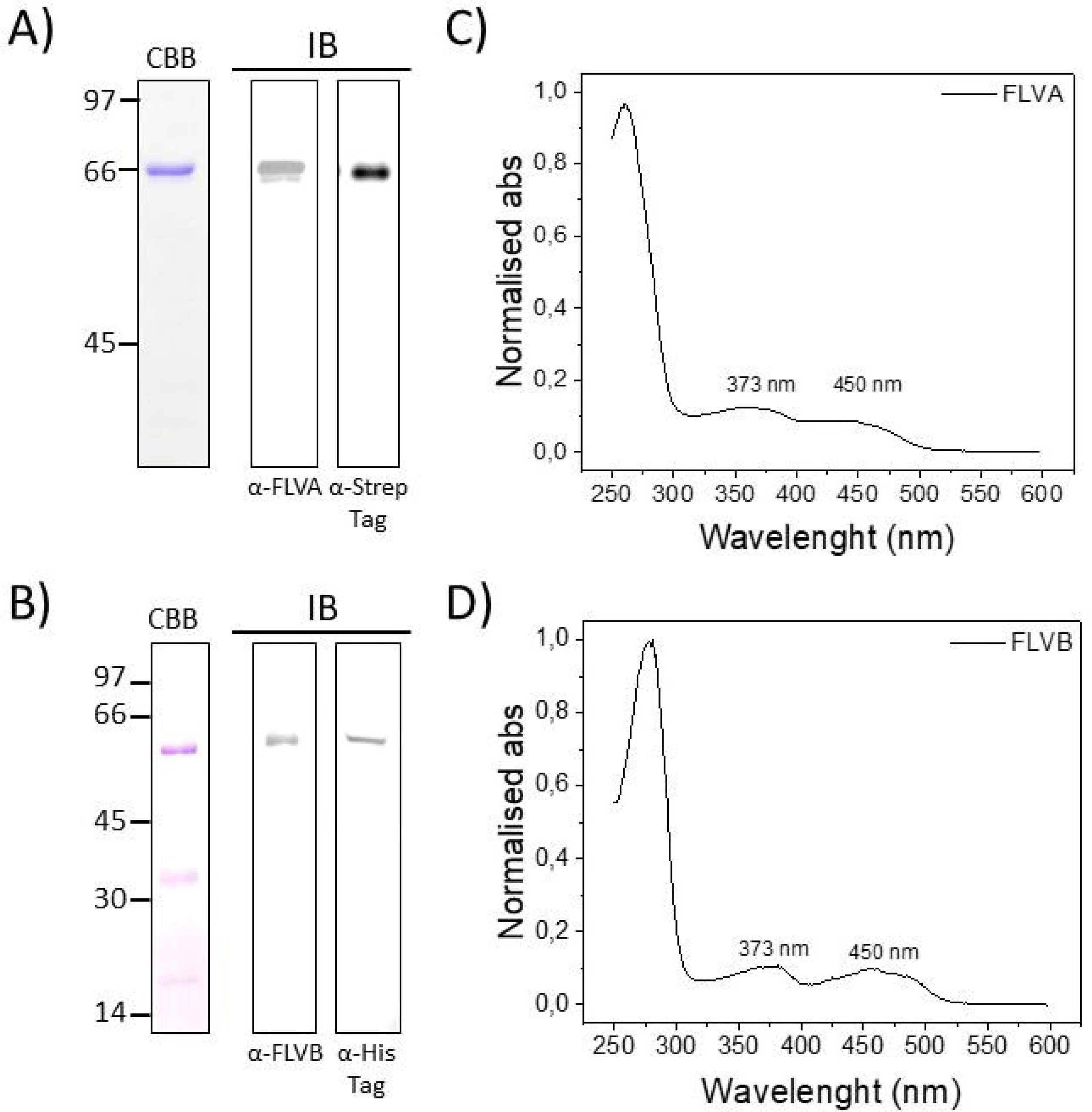
FLVA and FLVB biochemical properties. Coomassie Brilliant Blue-stained SDS-PAGE (CBB) of purified FLVA (A) and FLVB (B) and immunoblot analysis (IB) performed with antisera anti-FLVA and anti-Strep-Tag (A) and anti-FLVB and anti-His-Tag. On the left side, the apparent molecular weight of the ladder is reported in KDa. (B). UV-visible absorption spectra of purified FLVA (C) and FLVB (D). Absorption spectra of approximately 5 μM protein in a solution containing 30 mM Tris and 300 mM NaCl at pH 8.0 were recorded at room temperature.

The UV-vis absorption spectra of purified FLVA and FLVB displayed a primary peak at 280nm and two additional absorption peaks at 373nm and 450nm, characteristic of protein binding to the flavin moiety (Figure 2C-D) demonstrating that FMN was bound to the purified protein. Quantification of FMN content by the analysis of the absorption spectra estimated that each monomer of FLVA and FLVB bound approximately one molecule of flavin per monomer (1.09±0.26 for FLVA and 0.77±0.07 for FLVB). The presence of iron in purified FLVA and FLVB was verified via a colorimetric ferrozine-based assay that demonstrated that both proteins bound iron.

### *P. patens* FLVs are active and catalyze an NAD(P)H-dependent oxygen reduction reaction

Like all photosynthetic FLVs, *P. patens* FLVA and FLVB are characterized by a flavin reductase-like domain. To assess the activity of purified enzymes, we performed *in vitro* assays previously used to study other members of FDP enzyme family (17, 29). In particular, we conducted a spectroscopic assay to explore the ability of recombinant FLVA and FLVB to oxidize NADH and NADPH (Figure 3), that was already shown to be effective for cyanobacterial FLVs (16, 17). NADH oxidation, monitored from OD_340nm_, started as soon as FLVA and FLVB were added to the reaction solution, demonstrating the isolated proteins were active (Figure 3A). FLVB oxidized NADH faster than FLVA (Figure 3A) when the two enzymes were used at the same concentration, and 150μM NADH was fully oxidized by FLVB within a few minutes. The two enzymes were also able to oxidize NADPH (Figure 3B). Interestingly, FLVB confirmed higher efficiency compared to FLVA but, while FLVA displayed similar activity in presence of NADH and NADPH, FLVB showed a much higher rate with NADH.

**Figure 3.**
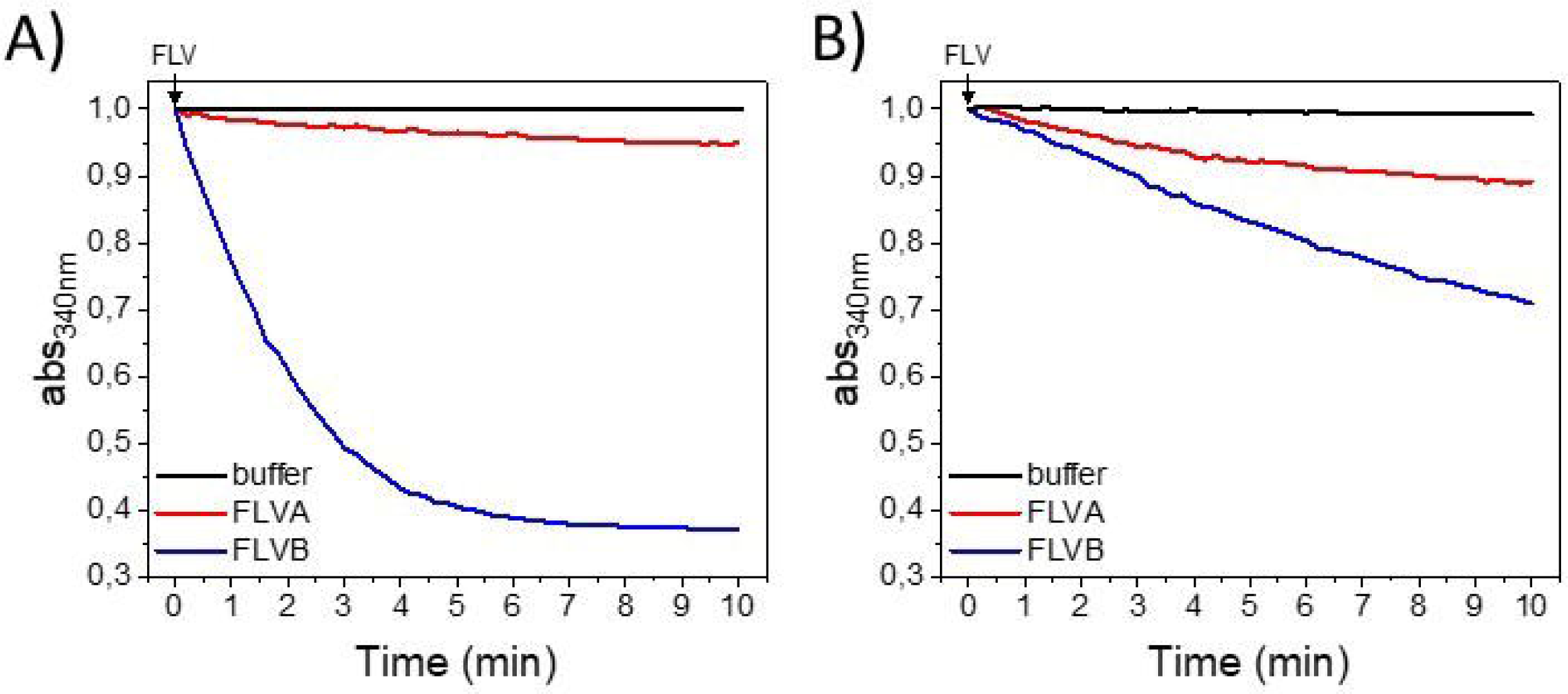
Kinetics of FLVA and FLVB NAD(P)H oxidase activity. FLVA and FLVB activity was tested in presence of 150μM NADH (A) and NADPH (B). Red lines correspond to 7μM FLVA and blue lines correspond to 7μM FLVB. Black lines correspond to 150μM NADH (A) and 150μM NADPH (B) without adding FLVs as negative controls. Traces were shifted to OD_340nm_ equal to 1 as starting point immediately after protein addition. Absorption spectra were recorded at room temperature.

In photosynthetic cells, FLVs perform O_2_ reduction to water. To independently confirm activity of purified proteins, we next assessed the capacity of recombinant *P. patens* FLVB to reduce O_2_ using a high-resolution respirometer (Figure 4). When FLVB was incubated in the measuring chamber, O_2_ levels were stable until NADH was added, which caused a fast decrease in O_2_ concentration, demonstrating that the protein indeed consumed O_2_. When O_2_ levels stabilized, a further addition of NADH resumed O_2_ reduction reaction, indicating that the enzyme was stable over time and reaction stopped because of exhaustion of reducing power (Figure 4). It is interesting to notice that O_2_ consumption was FLV-dependent, because there was no effect of NADH addition to the measuring chamber simply containing measuring buffer (Figure 4).

**Figure 4.**
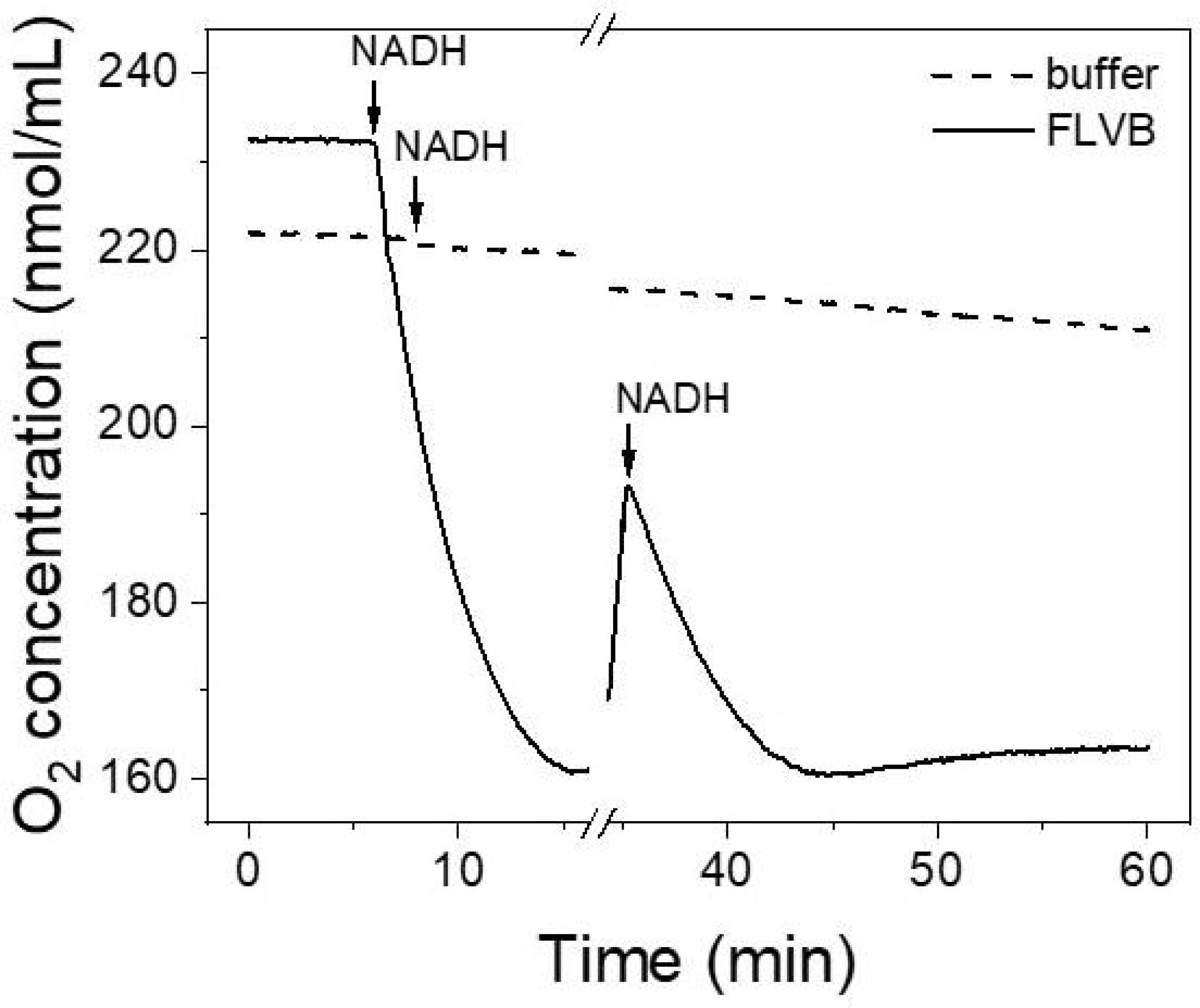
High-resolution respirometry of *P. patens* FLVB performed in an Oroboros Oxygraph-2k. The measurement started with sample buffer (Tris 30mM and NaCl 300mM) either containing 6µM FLVB (solid line) or not (dashed line). After about 7.5 minutes 150µM NADH was added to both samples. To verify protein stability, 70µM NADH was added again after about 35 minutes.

### FLVA and FLVB form stably bound and functional heterocomplex

FLVA and FLVB accumulation in *P. patens* needs the mutual presence of the two proteins since the depletion of one causes the destabilization of the other and the complete loss of FLV activity (12). Even though data suggest that FLVs are active together *in vivo*, an FLV heterocomplex has ever been purified and characterized so far, not even in cyanobacteria. To this aim, bacteria co-expressing 6XHisTag-FLVB and StrepIITag-FLVA were used to purify FLVA/FLVB complexes by two subsequent steps of immobilized metal affinity and Strep-Trap chromatography (Figure 5A). SDS-page showed the presence of two bands in the fractions eluted from the second purification column (Figure 5B and Figure S4). Immunoblot analysis confirmed the presence of both FLVA and FLVB that were successfully co-purified (Figure 5B and Figure S4). Coomassie staining revealed a similar signal for the two proteins, suggesting a stochiometric ratio between FLVA and FLVB (Figure 5B). This demonstrates that FLVA and FLVB indeed form a heterocomplex stable enough to survive purification.

**Figure 5.**
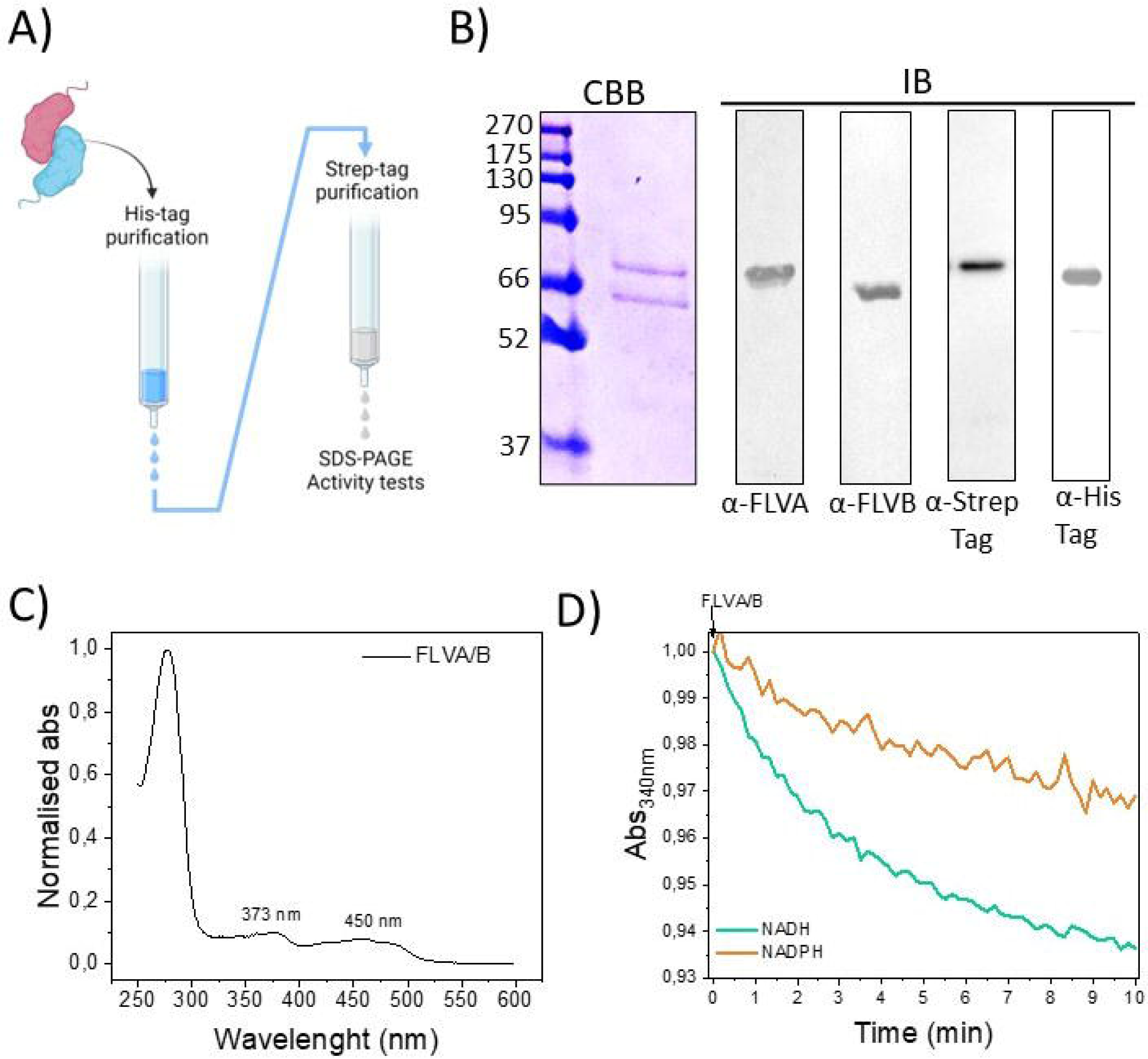
FLVA/FLVB heterocomplex properties. A) Bacteria co-expressing 6XHisTag-FLVB and StrepIITag-FLVA were purified by two subsequent steps of immobilized metal affinity and Strep-Trap chromatography. Purified FLVA/B heterocomplex was subjected to biochemical and functional analysis. B) Representative Coomassie Brilliant Blue-stained SDS-polyacrylamide gel and Western blot analysis of purified heterocomplex using α-FLVA and α-FLVB, α-StrepTag, α-HisTag antibodies. On the left side, the apparent molecular weight of the ladder is reported in KDa. C) Absorption spectrum of approximately 5 μM FLVA/B in a solution containing 30 mM Tris and 300 mM NaCl at pH 8.0 was recorded at room temperature. D) NAD(P)H oxidase activity of 1μM FLVA/B heterocomplex in presence of 150 μM NADH (green line) and 150 μM NADPH (orange line). Traces were shifted to OD_340nm_ equal to 1 as starting point immediately after protein adding. Absorption spectra were recorded at room temperature.

UV visible absorption spectrum of FLVA/B preparations showed the presence of flavin typical peaks (373nm, 450nm) that were present also in FLVA and FLVB when expressed separately (Figure 5C). About 1 molecule of FMN per FLV monomer (0.74±0.28) was associated to FLVA/FLVB complexes and spectroscopic analysis assessed the presence of iron ions. The FLVA/B heterocomplex was spectroscopically monitored for its ability to oxidize NADH and NADPH, exhibiting higher efficiency in the presence of NADH than NADPH (Figure 5D).

### Flavodiiron proteins complex formation require intermolecular disulfides bonds

Mature FLVA sequence and FLVB sequence contain respectively 16 and 9 cysteine residues that are compatible with redox sensing (8, 30) and may impact protein activity as it happens for other stromal enzymes like those of the Calvin-Benson Cycle (31). We examined the influence of the redox state on recombinant FLVA/B complexes by subjecting them to treatment with or without DTT as a reducing agent prior to SDS-PAGE analysis. In the absence of DTT, FLVA/B formed heterocomplexes that remained intact during separation on SDS-PAGE (Figure 6A). Exposure to DTT prompted the dissociation of FLVA/B heterocomplexes into monomers (Figure 6A), indicating the presence of inter-molecular disulfide bridges crucial for complex formation. Interestingly, both FLVA and FLVB exhibited similar behavior, forming homocomplexes under oxidizing conditions (Figure S5).

**Figure 6.**
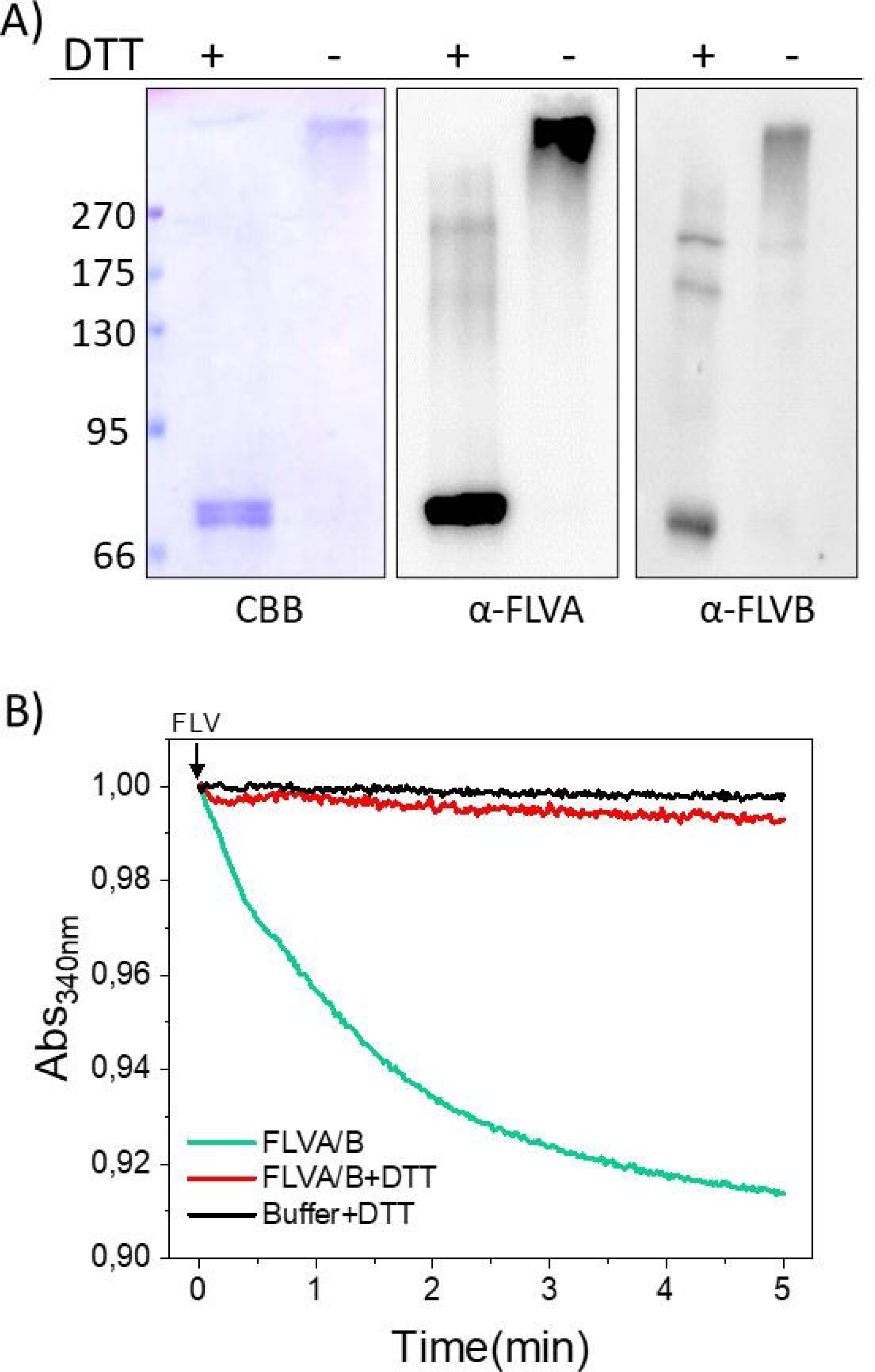
Redox dependence of flavodiiron proteins heterocomplex. A) SDS-PAGE and Western blotting of purified FLVA/B heterocomplex in presence or absence of reducing agent. On the left side, the apparent molecular weight of the ladder is reported in KDa. The samples were incubated at room temperature in the presence or absence of DTT as indicated on the top of each lane. Prior to SDS-page all samples were heated at 100^°^C. B) FLVA/B heterocomplex NADH oxidase activity in presence (red line) or absence (green line) of reducing agent. 4µM of purified enzyme and the equivalent volume of protein buffer (black line) were incubated at room temperature for ∼1hr with DTT prior to activity test. 150μM NADH was incubated with an equal volume of buffer and DTT and monitored as control (black line). Traces were shifted to OD_340nm_ equal to 1 as starting point immediately after protein adding. Absorption spectra were recorded at room temperature.

We next investigated enzymatic activity of purified FLVA/B when exposed to reducing conditions. Purified FLVs were incubated with DTT for 1 hour before testing their catalytic activity. Interestingly we observed a severe decrease in the enzyme efficiency suggesting that FLVs have activity is drastically altered in reducing conditions (Figure 6B). An equivalent volume of buffer was also incubated with the same concentration of DTT to exclude any potential interference of DTT on NADH absorbance.

## Discussion

### Expression of active recombinant eukaryotic FLVs including all cofactors

While various studies have explored the functional role of FLVs in living organisms (12, 26, 27), there is a lack of information about these proteins architecture and biochemical properties. To address this limitation, the present work investigated for the first time the biochemical and structural features of flavodiiron proteins of a eukaryotic photosynthetic organism. *P. patens* FLVA, FLVB and FLVA/B heterocomplexes expressed and purified from *E. coli* showed the expected molecular weight and bound flavin and iron atoms, as expected from the sequence analysis.

Notably, this is the case also for FLVA that exhibits a peculiar amino acid compositions in the diiron pocket domain that raised some doubts on its ability to coordinate iron (22). The analysis of *Synechocystis* sp. PCC6803 Flv1, an orthologue of FLVA, presented conflicting information. While the partial crystal structure of Flv1 revealed no iron (20), the full-length recombinant Flv1 was found to bind iron (17).

The iron coordination sites in the FLVB sequence revealed a type 1 canonical binding pattern characterized by Asp/Glu/His-rich residues (32). In contrast, FLVA, categorized as a type 2 protein, where the canonical diiron binding sites are substituted with neutral and basic residues (33). The data on *P. patens* FLVA suggest that despite possessing a non-canonical binding site (Type 2), these proteins are fully capable of coordinating iron. The type 2 binding site is a common occurrence in FDP in photosynthetic organisms, indicating that having two isoforms with distinct properties in the iron binding site is likely a conserved feature in FLV among photosynthetic organisms. The distinct properties of the iron binding sites may correlate with the observed higher activity in FLVB compared to FLVA. However, it should be noted that the characterized recombinant proteins expressed in bacteria might be influenced by the folding efficiency of the two proteins in the heterologous system.

### FLVs forms heterocomplexes stabilized by disulfide bridges

In the genomes of photosynthetic organisms, FLVs function as active pairs. In cyanobacteria, algae, and *P. patens*, the depletion of one protein results in the destabilization of the other, leading to the loss of all *in vivo* activity (7, 10, 12). In cyanobacteria, the overexpression of Flv3 in a *Δflv1* mutant background led to protein accumulation but proved insufficient for a complete functional complementation (24). Furthermore, recent studies showed that *P. patens* FLVs are functional in angiosperms when the two enzymes are expressed together (26, 27). In this study, we successfully isolated stable recombinant FLVA/B heterocomplexes using orthogonal purification and two different affinity tags. This demonstrated the strong interaction and ability of the two enzymes to form a heterocomplex. The FLVA/B complex exhibited enzymatic functionality in NAD(P)H oxidase activity. Notably, the overall activity of the heterocomplex was not higher than that of FLVB expressed alone. The potential advantage of having two distinct proteins is therefore likely not to enhance activity but to contribute to other features, such as regulation. It is noteworthy that the FLVA/B heterocomplex is stabilized by intermolecular cysteine bridges that persist through protein denaturation but can be dissociated into monomers upon incubation in reducing conditions.

When expressed in *E. coli*, both single FLVA and FLVB form homocomplexes under oxidizing conditions, like the FLVA/B heterocomplex, even though *in vivo* they are not even stable if expressed alone. The apparent contrast with *in vivo* observations can be attributed to differences in translation control between prokaryotes and plant cells where homocomplexes are likely recognized and degraded. Nevertheless, this demonstrates that FLVA and FLVB are similar enough to enable the formation of a stable and active complex.

Regulation of FLV activity. *In vivo*, FLV activity exhibits remarkable modulation. In *P. patens* it was shown to serve as the major contributor to electron transport for a few seconds following an increase in light intensity but then its activity becomes undetectable after 20 seconds of illumination. These observations suggest the existence of a rapid and efficient mechanism for the modulation of FLV activity.

One possibility is that FLV activity is regulated by competition for electron donors downstream of PSI. In the event of a sudden increase in illumination and the accumulation of electrons at the PSI acceptor side, FLV becomes available to accept electrons, reducing water and preventing nonspecific and potentially damaging reactions. Conversely, when other pathways, such as carbon fixation, are active, they may outcompete FLV for electron donors, limiting its activity. In *C. reinhardtii*, indications of competition between FLVs and hydrogenases are consistent with this hypothesis (19). This hypothesis is also consistent with observations presented here, such as the lower activity of the FLVA/B heterocomplex compared to FLVB alone and the protein lower affinity for NADPH compared to NADH, which is expected to be less abundant in the chloroplast stroma. In both cases, the biochemical observations align with the idea that, since the FLV reaction represents an energy loss, its activity should not be maximized but present only when needed and minimized when other pathways are active. Additionally, in accordance with early findings on FDPs (34, 35), it has been suggested that four electrons are necessary for the complete reduction of O_2_ to water, thereby preventing the formation of potentially harmful reactive oxygen species intermediates. It is conceivable that NADPH and ferredoxins may work in tandem, each donating two electrons. The proposed mechanism for the O_2_ reduction reaction of *P. patens* flavodiiron proteins is illustrated in Figure 7A. FLV exhibits a modular structure with a metallo-β-lactamase domain, a flavodoxin-like domain, and a flavin reductase-like domain. This modular arrangement exemplifies FLV as a protein with structurally independent units fused into a single entity (16).

**Figure 7.**
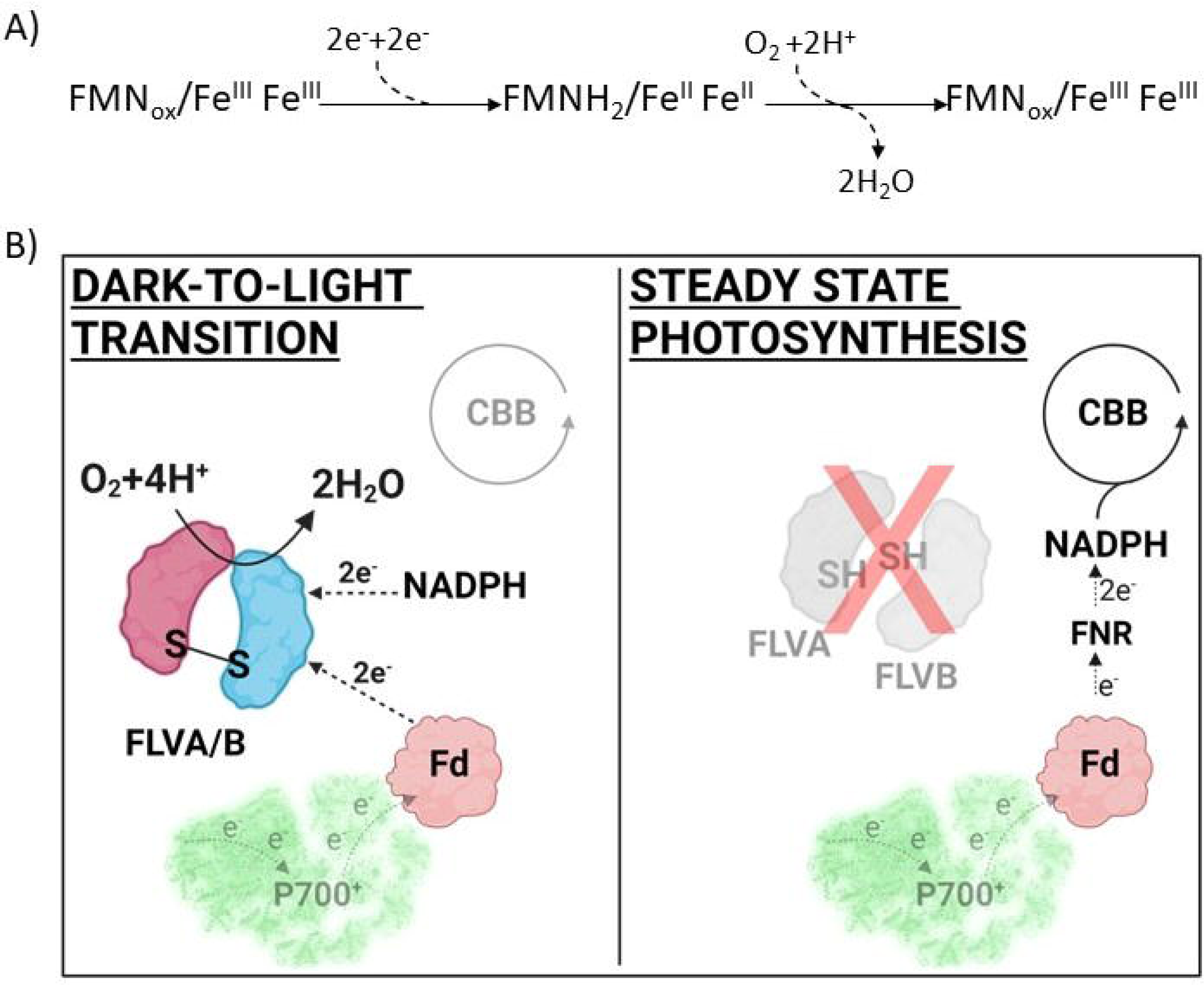
Proposed model describing *P. patens* flavodiiron protein catalytic activity. A) Overall proposed reaction scheme. Four electrons are necessary for the complete reduction of O_2_ to water, two electrons reduce the FMN moiety, while the other two the Fe-Fe centre. B) Redox dependence of FLV heterocomplex formation and activity. In oxidizing condition, in the dark-to-light transition, the disulfide bonds between monomer of FLVs are oxidized and the complex is formed and fully active. Electrons coming from Fd and NADPH participate to FLV activity. In reducing conditions, when photosynthesis is completely activated, disulfide bonds are cleaved, FLV complex is disassembled and decreases its catalytic efficiency.

Another plausible hypothesis, not necessarily mutually exclusive with substrate competition, is that FLV activity is regulated by the redox potential or indirectly by PSI and a change in ferredoxin affinity for its electron acceptors (18). Our findings indicate that upon reduction, FLV activity is strongly inhibited, possibly due to a direct effect or as an indirect consequence of heterocomplex disassembly. It is well-established that proteins in the stroma, particularly enzymes of the CBB cycle, are activated by a reducing potential under illumination. It would be particularly fitting if the same signal activating carbon fixation inhibits FLVs, a pathway competing for the same electrons. This regulatory mechanism would ensure active FLVs only when needed, i.e., when CO_2_ fixation is not fully active. Conversely, it would inhibit FLV activity when electrons could be utilized for growth instead of being wasted on oxygen reduction (Figure 7B).

## Experimental procedures

### FLVA and FLVB sequence and structural analysis

*P. patens* FLVA (UniProtKB A9RYP3, Gene PHYPA_018517) and FLVB (UniProtKB A9RQ00, Gene PHYPA_001217) sequences were aligned with MUSCLE alignment of MAFFT Multiple Sequence Alignment and secondary structures were predicted with JNet prediction tool. Results were visualized and analyzed with JalView. FLVA and FLVB structural domains were annotated from PROFILE database (PS50902) and from SMART database (SM00903). Model of FLVA and FLVB tertiary structures released by AlphaFold database (36) were also analysed and visualized with PyMOL software. To investigate residues and predicted structures conservation *Synechocystis sp. PCC6803* Flv1-ΔFlR (Uniprot P74373, PDB 60HC) was used as reference.

### FLV expression in Escherichia coli cells

*P. patens* FLVA and FLVB were expressed in their mature form, complete of the three domains typical of photosynthetic organisms. The N-terminal residues of putative chloroplast transit peptide were calculated with TargetP-2.0 (https://services.healthtech.dtu.dk/service.php?TargetP-2.0) and excluded from the expressed sequence. The CDS encoding FLVA (residues 55-723) was cloned into pETite® C-His Kan Vector (Lucigen) downstream of a sequence for a Strep-tag II (WSHPQFEK) to generate the recombinant plasmid pFLVA. The CDS encoding FLVB (residues 64-647) was cloned into pETDuet-1 (Novagen) downstream of a sequence for a hexa-histidine (6xHis) tag to generate the recombinant plasmid pFLVB. Primer sequences and a detailed cloning procedure are reported in Table S1. *E. coli T7 Shuffle* competent cells were transformed with pFLVA and pFLVB. For the heterocomplex purification, *E. coli T7 Shuffle* were simultaneously transformed with both plasmids and then plated on of Luria Bertani Agar medium (5g/L Yeast extract, 10g/L Tryptone, 3mL/1L NaOH 1M, 10g/L NaCl, Bacto agar 4g/L) containing specific antibiotics (50 μM kanamycin for FLVA, 100 μM ampicillin for FLVB and both kanamycin and ampicillin for FLVA/B). Colonies were pre-inoculated in 50mL of Luria Bertani (5g/L Yeast extract, 10g/L Tryptone, 3mL/1L NaOH 1M, 10g/L NaCl) liquid medium O/N and the pre-inoculum was then inoculated in 1 L of Terrific Broth (23,6g/L Yeast extract, 11,8g/L Tryptone, 9,4g/L K_2_HPO_4_, 2,2g/L KH_2_PO_4_, 4mL/L Glycerol liquid). Specific antibiotics were also added to the pre-inoculum and to the inoculum (kanamycin for FLVA, ampicillin for FLVB, kanamycin+ampicillin for FLVA/B). The growth rate was monitored by periodical OD_600nm_ measurements, riboflavin (Sigma-Aldrich) 50 μM and ammonium iron (II) sulfate (Mohr’s salt, (NH_4_)_2_Fe (SO_4_)_2_(H_2_O)_6_) 50 μM were added to the culture at OD_600nm_ ∼0.6 to promote their loading into proteins. 1 mM of isopropyl-β-D-thiogalactopyranoside (IPTG) was added at OD_600nm_ ∼0.8 to induce FLVA and/or FLVB expression. Bacterial liquid culture was then grown O/N at 16°C and 180 rpm. Cells were harvested by centrifugation at 20,000g for 20 min, at 4 °C.

### FLV purification from Escherichia coli cells

The harvested cells were resuspended in lysis buffer (30 mM Tris-HCl pH 8, 300mM NaCl and 0,1% V/V Triton x-100) and sonicated in the presence of lysozyme and protease inhibitors. After ultracentrifugation at 40,000g for 30 min, at 4 °C, the soluble fraction was collected for chromatography procedures. The soluble fraction was loaded onto Strep-Trap chromatography column (FLVA) (IBA-Lifesciences) or onto Immobilized Metal Affinity Chromatography column (FLVB) (Sigma-Aldrich) equilibrated with 20 mM Tris-HCl pH 8.0 and a salt gradient of 300mM NaCl. The fractions containing StrepIITag-FLVA or 6XHisTag-FLVB were eluted with 30mMTris and 300mM NaCl (pH 8.0). In case of FLVA/B heterocomplex purification, the bacterial soluble fraction was firstly loaded onto IMAC column, and the eluted material was then loaded into Strep-Trap chromatography column. Procedures and buffers were the same described above. Fractions containing FLVA, FLVB and FLVA/B were analyzed by SDS-page and then concentrated by ultra filtration (Sartorius, Vivaspin, 10 kDa cutoff). Analysis of different purification steps and quality of protein purifications was also assessed by Immunoblot analysis with custom-made α-FLVA, α-FLVB antibodies and commercial α-HisTag, α-StrepTag antibodies (Biorad). After transferring to nitrocellulose membrane, proteins were visualized with either Horseradish Peroxidase (HRP, Agrisera #AS09-60s) or Alkaline Phosphatase-conjugated secondary antibody (Sigma, #A3562). Notably, FLVB purification showed the highest purification yield (∼50μM FLVB/ 4ml elution volume) while FLVA and FLVA/B purification yield was 6 times lower (∼8μM FLVA/ 4ml elution volume).

### Determination of FLV biochemical properties

UV–visible absorption spectra of purified FLVs were recorded at room temperature with Agilent Cary 60 spectrophotometer. Proteins were diluted in Tris 30mM, Nacl 300mM, pH 8 solution before measurement. Protein concentration was determined considering the abs_280nm_ and ε=76320 M^-1^*cm^-1^ for FLVA, ε=66810 M^-1^*cm^-1^ for FLVB and ε=71565 M^-1^*cm^-1^ for FLVA/B. ε was determined from protein sequences using Expasy ProtParam tool. Flavin concentration of purified proteins was spectroscopically determined. Absorbance at 373nm and ε=12500 M^-1^*cm^-1^ was considered for quantitative FMN measurements. Iron content of purified FLV was also spectroscopically determined with a standard ferrozine assay (37). Briefly, 200 μL of each protein sample (5-10μM) were added with 50 μL of hydrochloric acid 12M, followed by heating the mixture for 20 minutes at 90°C. The mixture was centrifugated (16.000 g/15 minutes), and the resulting supernatant was treated with 60 μL ammonium acetate (NH_4_CH_3_CO_2_; oversaturated solution). After 10 minutes of incubation at room temperature, 10 μL ascorbic acid (C_6_H_8_O_6_; 75mM) were added, followed by other 10 minutes of incubation. Eventually, 50μL of FerroZine 10mM were added and after 30 minutes of incubation at room temperature, the absorption spectra of all samples were acquired. The presence of iron was estimated by measuring the absorbance between 450 and 750nm. Purified FLV displayed the typical iron absorption peak at 562nm, protein buffer was also used as negative control.

### Determination of FLV catalytic activity

FLV catalytic activity was determined by spectroscopic analysis. The assay mixture contained FLVs and NAD(P)H in Tris 30mM, NaCl 300mM, pH 8. NAD(P)H concentration was monitored by measuring absorption spectra (200-400nm) every 12 seconds to follow the decrease at 340nm at room temperature. For reducing conditions, purified FLV were pre-incubated for 1hour with DTT 50mM before NAD(P)H oxidation assay (38). An equivalent volume of buffer was incubated with DTT 50mM as negative control. All redox treatments were carried out in 30 mM Tris, 300Mm NaCl, pH 8.

FLV oxygen consumption rates were measured by a respirometer at 25°C (O2K respirometer Oroboros ®). Purified FLVB was used at final concentration of ∼6μM and incubated with ∼150μM NADH in Tris 30mM, NaCl 300mM (2ml final volume). Oxygen concentration level (nmol/mL) in the sealed chamber was recorded over time for 60 minutes at room temperature.

### Immunoblot analysis of purified proteins in oxidizing and reducing conditions

FLV enzymes were purified under non-reducing conditions allowing the formation of regulatory disulfide bonds. About 1μg of purified FLVA, FLVB, FLVA-FLVB were treated with sample buffer (Tris-HCl 125 Mm, pH 6.8; glycerol 30 % (w/v); sodium dodecyl sulphate 9 % (w/v); 0.04% (w/v) bromophenol blue). In reducing conditions dithiothreitol (100 mM) was also added to the sample buffer. Proteins were denatured at 100°C for 1 min, centrifugation at 13.000 g for 5 min and then loaded onto SDS-PAGE (10% or 12% acrylamide). Proteins were stained by Brilliant Blue Coomassie or transferred to nitro-cellulose membranes (Pall Corporation) for immunodetection with custom-made α-FLVA, α-FLVB antibodies as described above.

## Data availability

All data are contained within the manuscript.

## Supporting information

This article contains supporting information.

## Acknowledgments

We thank Stefano Liberi, Monica Chinellato, Elisa Barbazza and Lorenzo Iacobucci for FLV cloning and setting up FLV purification. The authors would like to acknowledge the use of ChatGPT3.5 to revise some part of the text. TM thanks Oroboros Instruments GmbH for their support with O2k-PhotoBiology.

## Author contributions

Conceptualization, A.A. and T.M.; Investigation, C.B., E.T., M.B.; Supervision, L.C., A.A., T.M.; Writing – original draft, A.A. and C.B.; Writing – review & editing, T.M, L.C.

## Funding and additional information

AA acknowledges funding from MUR PRIN2022PNRR - IRONCROP (P2022ZXWLK).

## References

1. Sonoike, K., and Terashima, I. (1994) Mechanism of photosystem-I photoinhibition in leaves ofCucumis sativus L. Planta. 194, 287–293

2. Tiwari, A., Mamedov, F., Grieco, M., Suorsa, M., Jajoo, A., Styring, S., Tikkanen, M., and Aro, E.-M. (2016) Photodamage of iron-sulphur clusters in photosystem I induces non-photochemical energy dissipation. Nat Plants. 2, 16035

3. Lima-Melo, Y., Gollan, P. J., Tikkanen, M., Silveira, J. A. G., and Aro, E.-M. (2019) Consequences of photosystem-I damage and repair on photosynthesis and carbon use in Arabidopsis thaliana. Plant J. 97, 1061–1072

4. Asada, K. (2000) The water-water cycle as alternative photon and electron sinks. Philos Trans R Soc Lond B Biol Sci. 355, 1419–1431

5. Walker, B. J., Strand, D. D., Kramer, D. M., and Cousins, A. B. (2014) The response of cyclic electron flow around photosystem I to changes in photorespiration and nitrate assimilation. Plant Physiol. 165, 453–462

6. Alboresi, A., Storti, M., and Morosinotto, T. (2019) Balancing protection and efficiency in the regulation of photosynthetic electron transport across plant evolution. New Phytol. 221, 105–109

7. Helman, Y., Tchernov, D., Reinhold, L., Shibata, M., Ogawa, T., Schwarz, R., Ohad, I., and Kaplan, A. (2003) Genes encoding A-type flavoproteins are essential for photoreduction of O2 in cyanobacteria. Curr Biol. 13, 230–235

8. Ermakova, M., Battchikova, N., Richaud, P., Leino, H., Kosourov, S., Isojärvi, J., Peltier, G., Flores, E., Cournac, L., Allahverdiyeva, Y., and Aro, E.-M. (2014) Heterocyst-specific flavodiiron protein Flv3B enables oxic diazotrophic growth of the filamentous cyanobacterium Anabaena sp. PCC 7120. Proc. Natl. Acad. Sci. U.S.A. 111, 11205–11210

9. Santana-Sanchez, A., Solymosi, D., Mustila, H., Bersanini, L., Aro, E.-M., and Allahverdiyeva, Y. (2019) Flavodiiron proteins 1-to-4 function in versatile combinations in O2 photoreduction in cyanobacteria. Elife. 8, e45766

10. Chaux, F., Burlacot, A., Mekhalfi, M., Auroy, P., Blangy, S., Richaud, P., and Peltier, G. (2017) Flavodiiron Proteins Promote Fast and Transient O2 Photoreduction in Chlamydomonas. Plant physiology. 10.1104/pp.17.00421

11. Shimakawa, G., Ishizaki, K., Tsukamoto, S., Tanaka, M., Sejima, T., and Miyake, C. (2017) The Liverwort, Marchantia, Drives Alternative Electron Flow Using a Flavodiiron Protein to Protect PSI. Plant Physiol. 173, 1636–1647

12. Gerotto, C., Alboresi, A., Meneghesso, A., Jokel, M., Suorsa, M., Aro, E.-M., and Morosinotto, T. (2016) Flavodiiron proteins act as safety valve for electrons in Physcomitrella patens. Proc Natl Acad Sci U S A. 113, 12322–12327

13. Bag, P., Shutova, T., Shevela, D., Lihavainen, J., Nanda, S., Ivanov, A. G., Messinger, J., and Jansson, S. (2023) Flavodiiron-mediated O2 photoreduction at photosystem I acceptor-side provides photoprotection to conifer thylakoids in early spring. Nat Commun. 14, 3210

14. Folgosa, F., Martins, M. C., and Teixeira, M. (2018) Diversity and complexity of flavodiiron NO/O2 reductases. FEMS Microbiol Lett. 10.1093/femsle/fnx267

15. Martins, M. C., Romão, C. V., Folgosa, F., Borges, P. T., Frazão, C., and Teixeira, M. (2019) How superoxide reductases and flavodiiron proteins combat oxidative stress in anaerobes. Free Radic Biol Med. 140, 36–60

16. Vicente, J. B., Gomes, C. M., Wasserfallen, A., and Teixeira, M. (2002) Module fusion in an A-type flavoprotein from the cyanobacterium Synechocystis condenses a multiple-component pathway in a single polypeptide chain. Biochem Biophys Res Commun. 294, 82–87

17. Brown, K. A., Guo, Z., Tokmina-Lukaszewska, M., Scott, L. W., Lubner, C. E., Smolinski, S., Mulder, D. W., Bothner, B., and King, P. W. (2019) The oxygen reduction reaction catalyzed by Synechocystis sp. PCC 6803 flavodiiron proteins. Sustainable Energy Fuels. 3, 3191–3200

18. Sétif, P., Shimakawa, G., Krieger-Liszkay, A., and Miyake, C. (2020) Identification of the electron donor to flavodiiron proteins in Synechocystis sp. PCC 6803 by in vivo spectroscopy. Biochim Biophys Acta Bioenerg. 1861, 148256

19. Jokel, M., Nagy, V., Tóth, S. Z., Kosourov, S., and Allahverdiyeva, Y. (2019) Elimination of the flavodiiron electron sink facilitates long-term H2 photoproduction in green algae. Biotechnol Biofuels. 12, 280

20. Borges, P. T., Romão, C. V., Saraiva, L. M., Gonçalves, V. L., Carrondo, M. A., Teixeira, M., and Frazão, C. (2019) Analysis of a new flavodiiron core structural arrangement in Flv1-ΔFlR protein from Synechocystis sp. PCC6803. Journal of Structural Biology. 205, 91–102

21. Allahverdiyeva, Y., Isojärvi, J., Zhang, P., and Aro, E.-M. (2015) Cyanobacterial Oxygenic Photosynthesis is Protected by Flavodiiron Proteins. Life (Basel). 5, 716–743

22. Alboresi, A., Storti, M., Cendron, L., and Morosinotto, T. (2019) Role and regulation of class-C flavodiiron proteins in photosynthetic organisms. Biochem J. 476, 2487–2498

23. Allahverdiyeva, Y., Ermakova, M., Eisenhut, M., Zhang, P., Richaud, P., Hagemann, M., Cournac, L., and Aro, E.-M. (2011) Interplay between Flavodiiron Proteins and Photorespiration in Synechocystis sp. PCC 6803. J Biol Chem. 286, 24007–24014

24. Allahverdiyeva, Y., Mustila, H., Ermakova, M., Bersanini, L., Richaud, P., Ajlani, G., Battchikova, N., Cournac, L., and Aro, E.-M. (2013) Flavodiiron proteins Flv1 and Flv3 enable cyanobacterial growth and photosynthesis under fluctuating light. Proc Natl Acad Sci U S A. 110, 4111–4116

25. Shimakawa, G., Shoguchi, E., Burlacot, A., Ifuku, K., Che, Y., Kumazawa, M., Tanaka, K., and Nakanishi, S. (2022) Coral symbionts evolved a functional polycistronic flavodiiron gene. Photosynth Res. 151, 113–124

26. Yamamoto, H., Takahashi, S., Badger, M. R., and Shikanai, T. (2016) Artificial remodelling of alternative electron flow by flavodiiron proteins in Arabidopsis. Nat Plants. 2, 16012

27. Wada, S., Yamamoto, H., Suzuki, Y., Yamori, W., Shikanai, T., and Makino, A. (2018) Flavodiiron Protein Substitutes for Cyclic Electron Flow without Competing CO2 Assimilation in Rice. Plant Physiol. 176, 1509–1518

28. Wasserfallen, A., Ragettli, S., Jouanneau, Y., and Leisinger, T. (1998) A family of flavoproteins in the domains Archaea and Bacteria. Eur J Biochem. 254, 325–332

29. Folgosa, F., Martins, M. C., and Teixeira, M. (2018) The multidomain flavodiiron protein from Clostridium difficile 630 is an NADH:oxygen oxidoreductase. Sci Rep. 8, 10164

30. Zhang, P., Eisenhut, M., Brandt, A.-M., Carmel, D., Silén, H. M., Vass, I., Allahverdiyeva, Y., Salminen, T. A., and Aro, E.-M. (2012) Operon flv4-flv2 provides cyanobacterial photosystem II with flexibility of electron transfer. Plant Cell. 24, 1952–1971

31. Gurrieri, L., Fermani, S., Zaffagnini, M., Sparla, F., and Trost, P. (2021) Calvin–Benson cycle regulation is getting complex. Trends in Plant Science. 26, 898–912

32. Romão, C. V., Vicente, J. B., Borges, P. T., Victor, B. L., Lamosa, P., Silva, E., Pereira, L., Bandeiras, T. M., Soares, C. M., Carrondo, M. A., Turner, D., Teixeira, M., and Frazão, C. (2016) Structure of Escherichia coli Flavodiiron Nitric Oxide Reductase. Journal of Molecular Biology. 428, 4686–4707

33. Gonçalves, V. L., Saraiva, L. M., and Teixeira, M. (2011) Gene expression study of the flavodiiron proteins from the cyanobacterium Synechocystis sp. PCC6803. Biochem Soc Trans. 39, 216–218

34. Di Matteo, A., Scandurra, F. M., Testa, F., Forte, E., Sarti, P., Brunori, M., and Giuffrè, A. (2008) The O2-scavenging flavodiiron protein in the human parasite Giardia intestinalis. J Biol Chem. 283, 4061–4068

35. Frederick, R. E., Caranto, J. D., Masitas, C. A., Gebhardt, L. L., MacGowan, C. E., Limberger, R. J., and Kurtz, D. M. (2015) Dioxygen and nitric oxide scavenging by Treponema denticola flavodiiron protein: a mechanistic paradigm for catalysis. J Biol Inorg Chem. 20, 603–613

36. Jumper, J., Evans, R., Pritzel, A., Green, T., Figurnov, M., Ronneberger, O., Tunyasuvunakool, K., Bates, R., Žídek, A., Potapenko, A., Bridgland, A., Meyer, C., Kohl, S. A. A., Ballard, A. J., Cowie, A., Romera-Paredes, B., Nikolov, S., Jain, R., Adler, J., Back, T., Petersen, S., Reiman, D., Clancy, E., Zielinski, M., Steinegger, M., Pacholska, M., Berghammer, T., Bodenstein, S., Silver, D., Vinyals, O., Senior, A. W., Kavukcuoglu, K., Kohli, P., and Hassabis, D. (2021) Highly accurate protein structure prediction with AlphaFold. Nature. 596, 583–589

37. Stookey, L. L. (1970) Ferrozine---a new spectrophotometric reagent for iron. Anal. Chem. 42, 779–781

38. Cardi, M., Zaffagnini, M., De Lillo, A., Castiglia, D., Chibani, K., Gualberto, J. M., Rouhier, N., Jacquot, J.-P., and Esposito, S. (2016) Plastidic P2 glucose-6P dehydrogenase from poplar is modulated by thioredoxin m-type: Distinct roles of cysteine residues in redox regulation and NADPH inhibition. Plant Science. 252, 257–266

