## Supplemental Figures and Table for "*Physcomitrium patens* flavodiiron proteins form a redox-dependent heterocomplex"

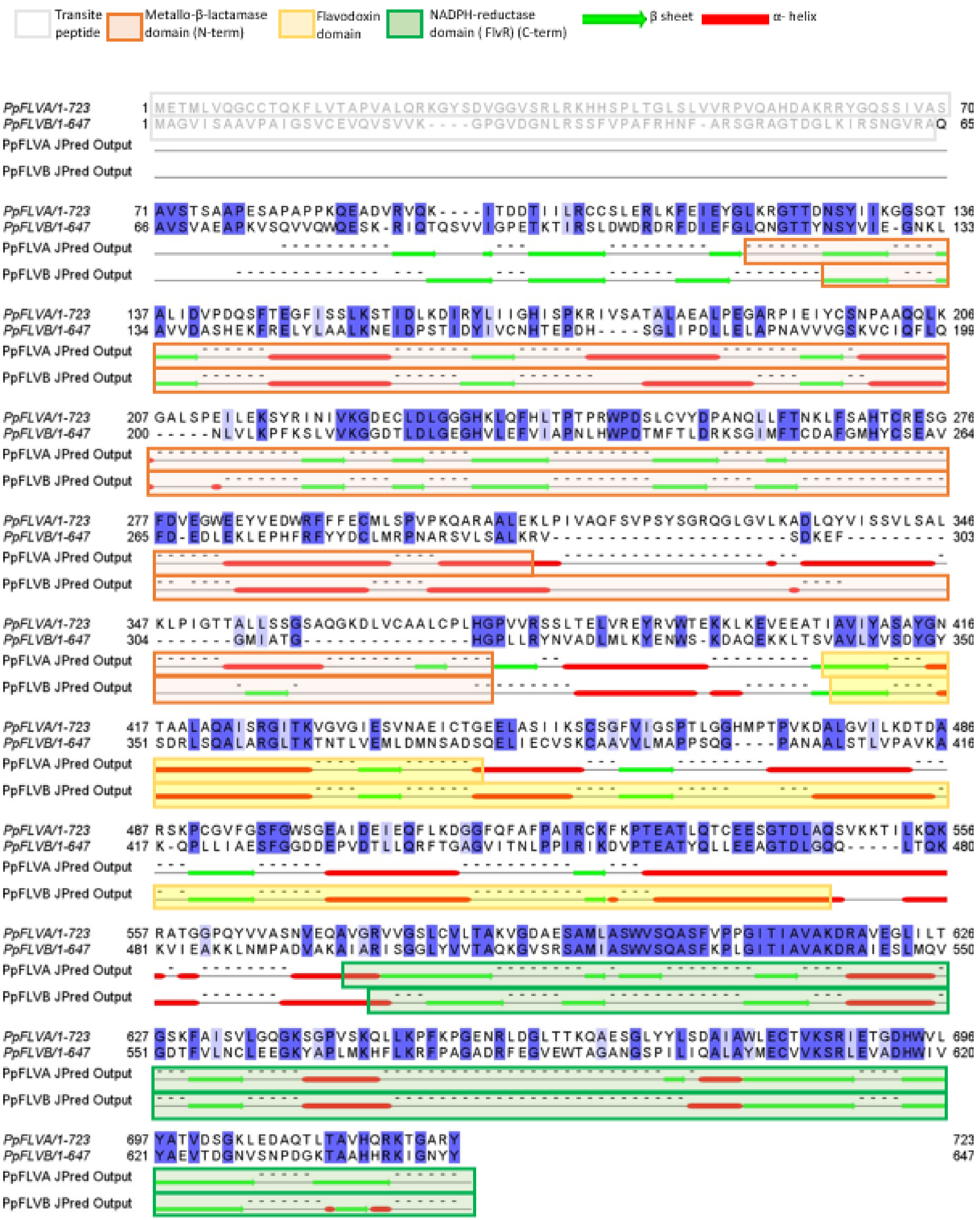

**Figure S1. Sequence and structural domain analysis of *Physcomitrium patens* FLVA and FLVB.** The protein sequence alignment generated using MUSCLE, and the predicted secondary structures from JNET for *P. patens* FLVA and FLVB were visualized using Jalview, following manual adjustments. Residues were color-coded based on percentage identity (>80% in dark blue, between 60% and 80% in light blue). Predicted α-helices and β-sheets were indicated by green arrows and red boxes, respectively. The metallo-β-lactamase domain, annotated from the SMART database (SM00849) was labeled in orange, the flavodoxin-like domain from the PROFILE database (PS50902) was labeled in yellow, and the flavin reductase-like domain from the SMART database (SM00903) was labeled in green. The predicted cleavage sites for FLVA and FLVB were positioned at amino acid 70 and 65 and the transit peptide was labeled in grey.

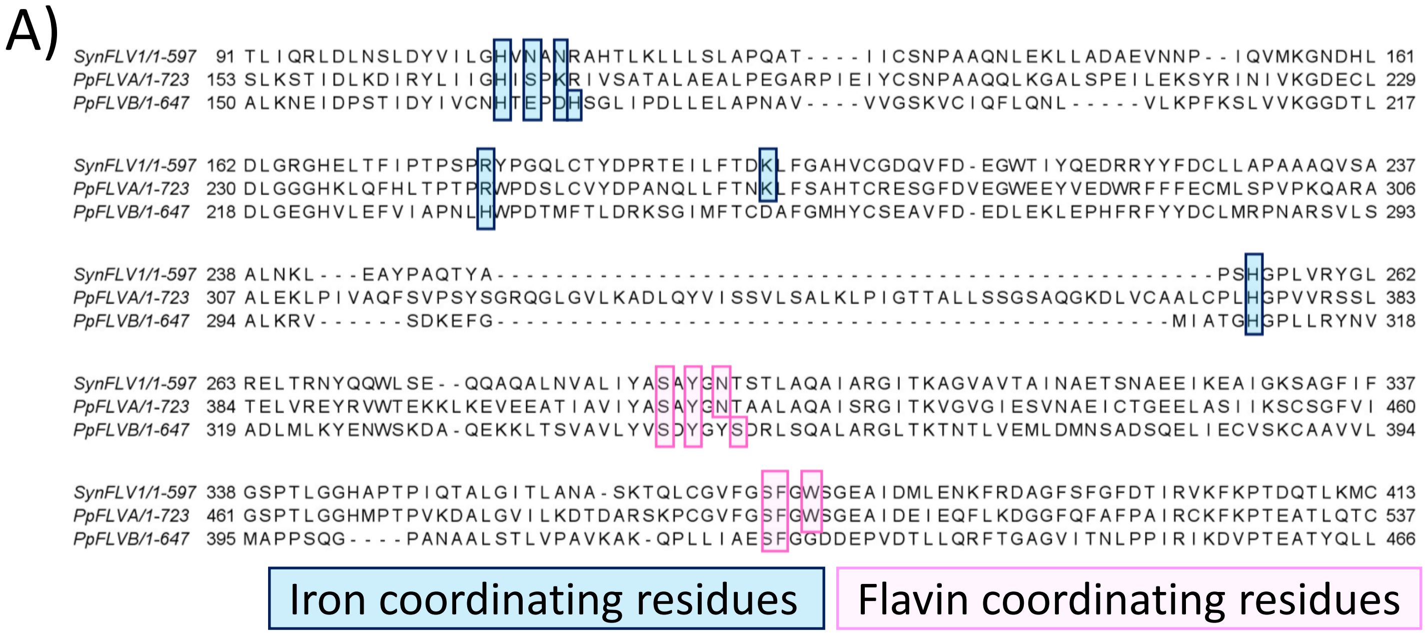


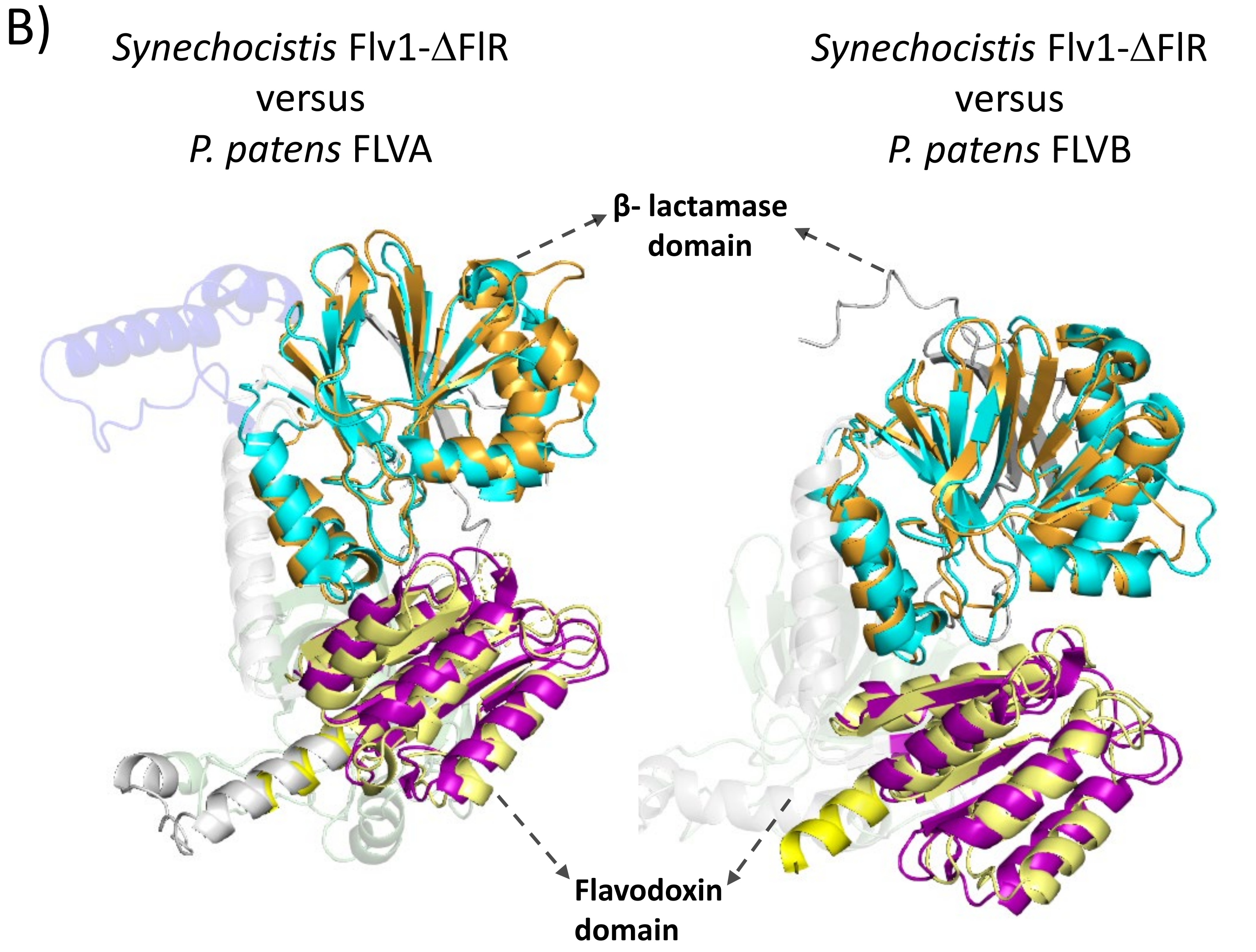


**Figure S2. Structural and sequence comparison between *P. patens* FLV and *Synechocistis* Flv1-DFlR.** A) Alignment of *Synechocystis* Flv1 and *P. patens* FLVA and PpFLVB. Residues predicted to coordinate flavin are highlighted in pink and those coordinating iron are highlighted in blue squares. B) Structural superposition of *Synechocistis* Flv1-ΔFlR protein and AlphaFold predictions of *P. patens* FLVA (left panel) and FLVB (right panel). The β-lactamase (residues 59-254) and flavodoxin-like domains (residues 283-421) from Flv1-ΔFlR are respectively coloured in blue and purple, while those from *P. patens* FLVA and FLVB are coloured respectively in orange and yellow. The remaining parts of *P. patens* FLVA and FLVB are reported as transparent domains.

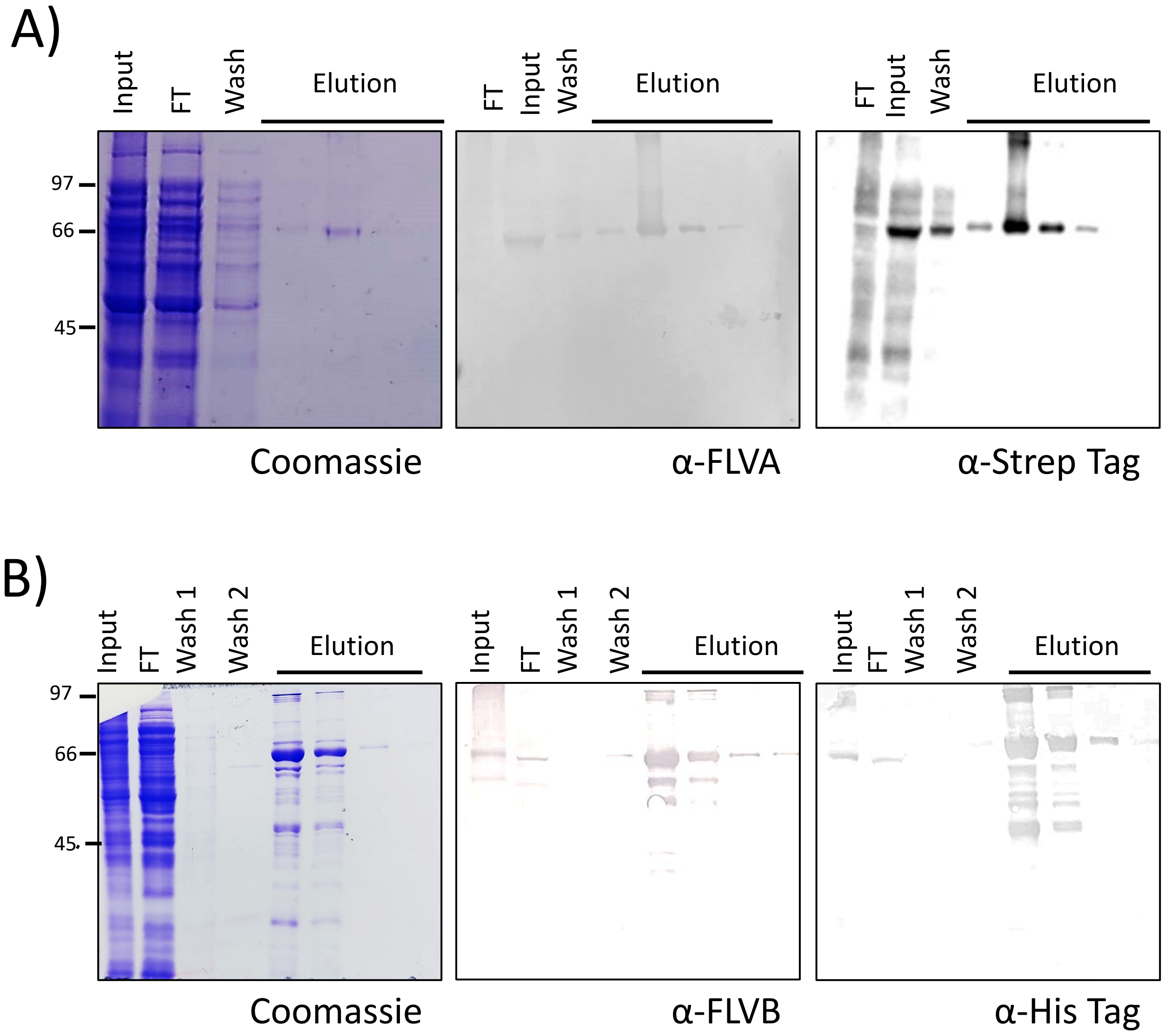


**Figure S3. Purification of and StrepIITag-FLVA (A) and 6XHisTag-FLVB (B) from the soluble fractions of *Escherichia coli* total protein extracts.** Coomassie Brilliant Blue-stained SDS-polyacrylamide gel with fractions obtained during Strep-tag (A) and His-tag (B) affinity chromatography purification of StrepTagII-FLVA and 6XHisTagFLVB. Immunoblot analysis was performed using α-FLVA, α-StrepTag, α-FLVB and α-His Tag antibodies. Input: total soluble proteins; FT: flow through; wash: proteins harvested after 20mM Tris-300mM NaCl; wash 1: proteins harvested after 20mM Tris-300mM NaCl; wash 2: proteins harvested after 5mM imidazole; Elution after 4.5mM desthiobiotin (A), Elution after 250mM imidazole (B).

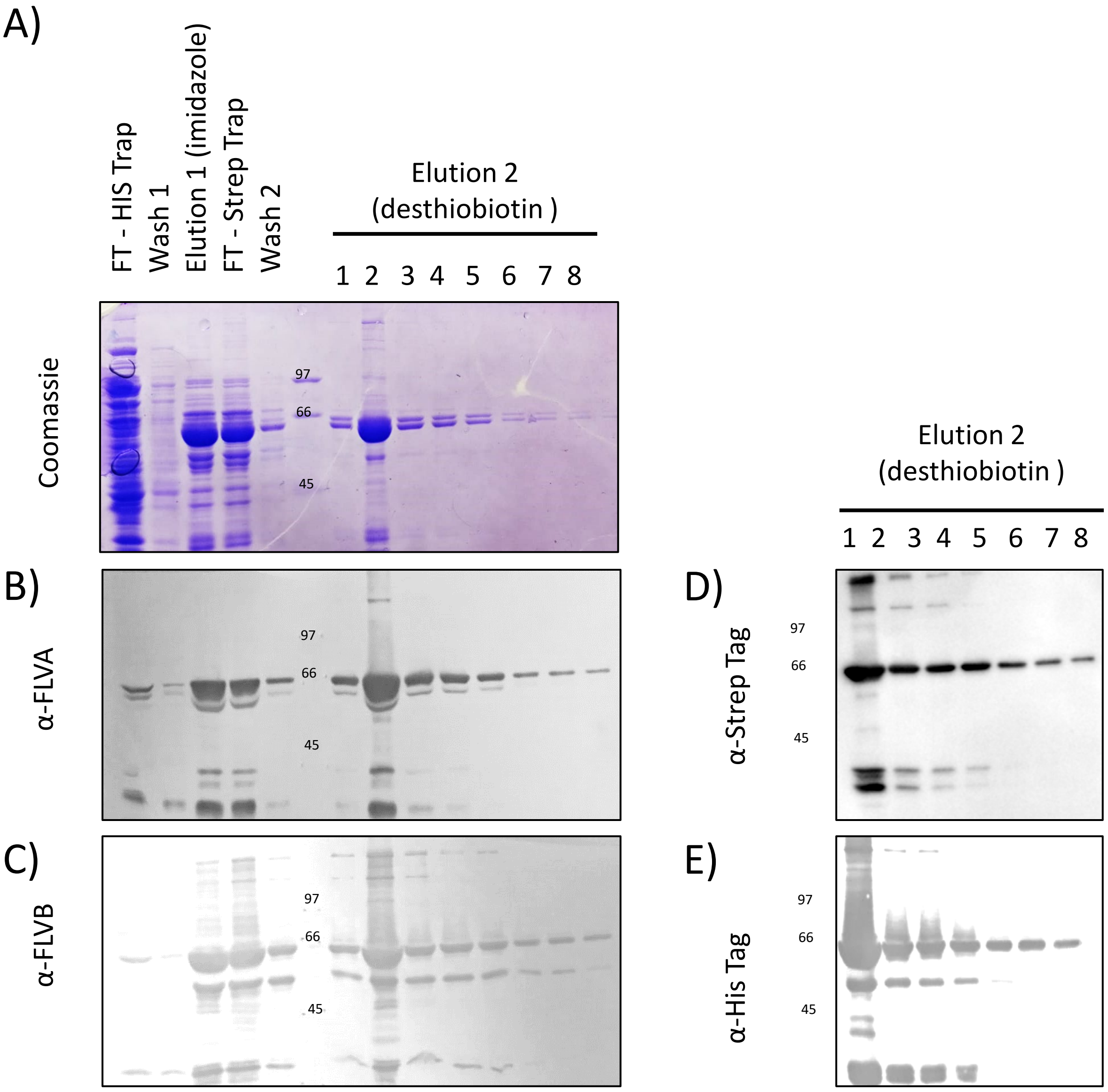


**Figure S4. Purification of 6XHisTagFLVB and-StrepIITag-FLVA heterocomplex from the soluble fractions.** A) Coomassie Brilliant Blue-stained SDS-polyacrylamide gel with fractions obtained during immobilized metal affinity chromatography (IMAC) coupled with Strep-Tag affinity chromatography purification of recombinant 6XHisTagFLVB and-StrepIITag-FLVA heterocomplex . Lane 1 HisTrap column flow through, lane 2, after wash with 20 mM imidazole buffer; lane 3, after elution using 250 mM imidazole buffer; lane 4, Strep-trap flow through, lane 5, Strep-trap wash, lane 6a-6g, 4.5mM desthiobiotin elution, Molecular weight markers are also indicated (in thousands). B-C) Western blot analysis of the different purification steps using α-FLVB, α-FLVA antibodies. D-E) Western blot analysis of the elute with 4.5mM desthiobiotin buffer using α-His Tag, α-StrepTag antibodies.

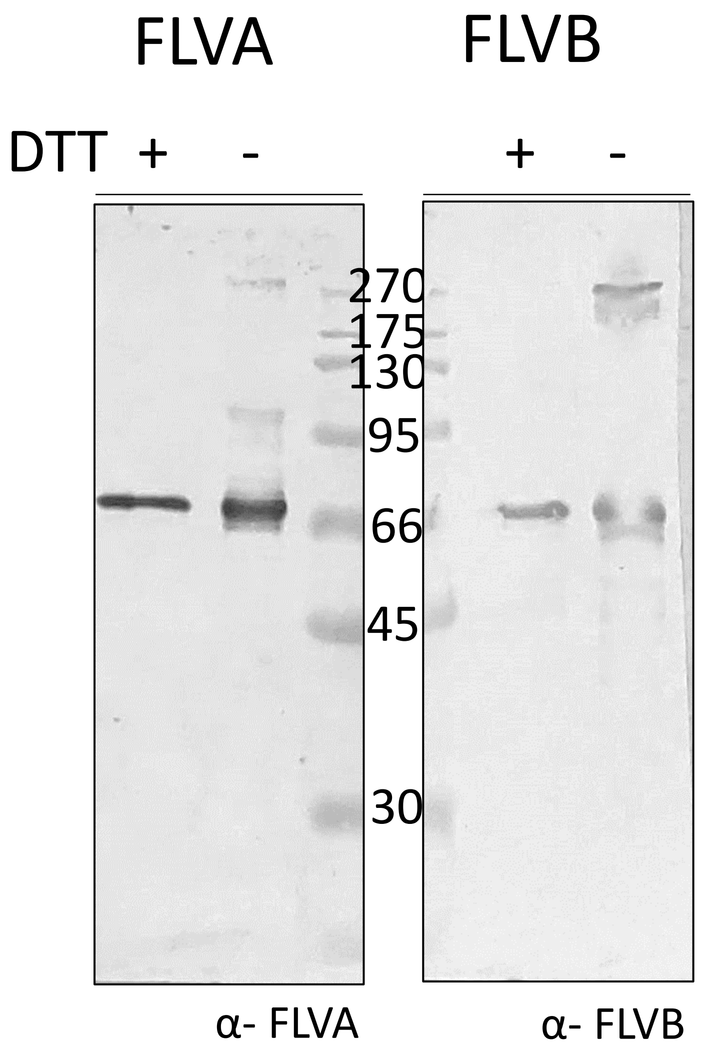


**Figure S5. Redox dependence of flavodiiron proteins homocomplexes.** SDS-PAGE and Western blotting of purified FLVA and FLVB homocomplex in presence or absence of reducing agent. The samples were incubated at room temperature in the presence (33 mM) or absence of DTT as indicated on the top of each lane. Prior to SDS-page all samples were heated at 100˚ . MW standards (in thousands).

| Primer name | Gene | Sequence | Use (RE) |
|---|---|---|---|
| FLVA_For_TAG | FLVA | GAAGGAGATATACATatgGCGAGCGCGTGGAGCCACCCGCAGTTCGAAAAAgctcatgatgcgaaaagaagatatgg | pFLVA vector |
| FLVA_Rev | FLVA | GTGATGGTGGTGATGATGttaGTAACGAGCACCCGTCTTGCg | pFLVA vector |
| FLVB_For_TAG | FLVB | CAGGATCCGGCGCAGGCTGTGAGTGTGG | pFLVB vector (BamHI) |
| FLVB_Rev | FLVB | GGCGCGCCTCAATAGTAGTTCCCAAT | pFLVB vector (AscI) |

**Table S1. Primers employed for molecular cloning and screening.** The table reports all the primers used to generate vectors for the expression of *Physcomitrium patens* FLVA and FLVB. Primers were designed to clone FLVA coding sequence in frame with Strep-II-Tag in pETite® C-His Kan Vector and and FLVB coding sequence in frame with 6XHis-Tag in pETDuet-1 vectors. cDNA encoding PpFLVA and PpFLVB was amplified by PCR using respectively FLVA_For_TAG/FLVA_Rev and FLVB_For_TAG/FLVB_Rev primers and the cDNA encoding PpFlvA-2A-PpFlvB as template (Yamamoto et al., 2016). FLVA PCR product was cloned into pETite® C-His Kan Vector with enzyme-free cloning (Expresso™ T7 Cloning and Expression System). FLVB PCR product were first cloned in pJET1.2/blunt cloning vector (Thermofisher) and then digested with BamHI and AscI for cloning into pETDuet-1 vectors.
